# Hepatitis B Virus Infection as a Neglected Tropical Disease

**DOI:** 10.1101/164491

**Authors:** Geraldine A O’Hara, Anna L McNaughton, Tongai Maponga, Pieter Jooste, Ponsiano Ocama, Roma Chilengi, Jolynne Mokaya, Mitchell I Liyayi, Tabitha Wachira, David M Gikungi, Lela Burbridge, Denise O’Donnell, Connie S Akiror, Derek Sloan, Judith Torimiro, Louis Marie Yindom, Robert Walton, Monique Andersson, Kevin Marsh, Robert Newton, Philippa C Matthews

## BACKGROUND

The Global Hepatitis Health Sector Strategy is aiming for ‘elimination of viral hepatitis as a public health threat’ by 2030 [1], while enhanced elimination efforts for hepatitis are also promoted under the broader remit of global Sustainable Development Goals (SDGs) [2]. This is an enormous challenge for hepatitis B virus (HBV) given the estimated global burden of 260 million chronic carriers, of whom the majority are unaware of their infection [3] (Figure 1).

**Figure 1.**
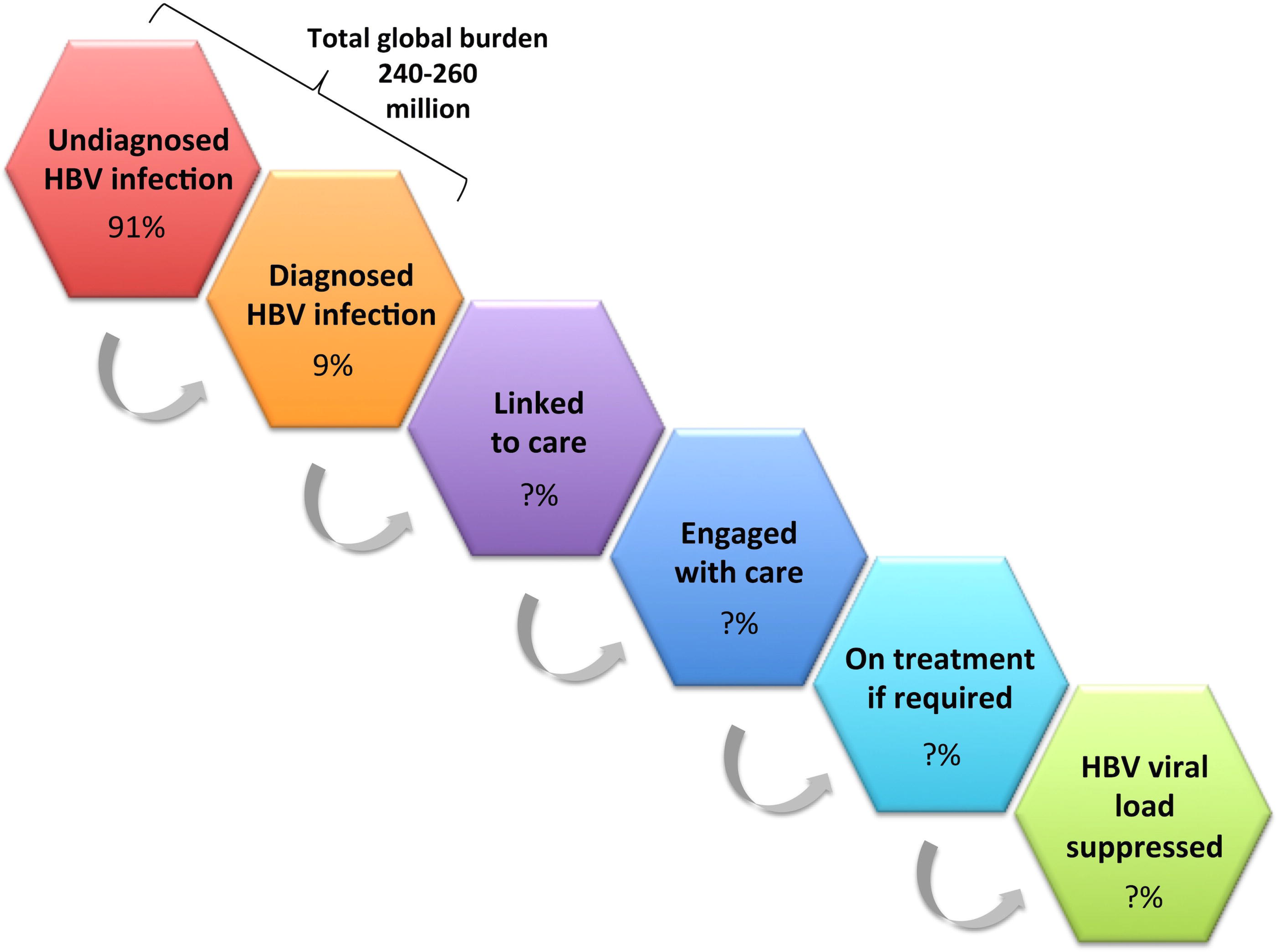
The HBV cascade. Diagrammatic representation of the total burden of HBV infection, and the subsets of individuals who are diagnosed (orange), linked to care (purple), engaged with care (dark blue), on treatment (light blue) and have suppressed viraemia (green). An estimate of the proportion of cases undiagnosed vs. diagnosed (91% vs. 9%, respectively) is based on the WHO factsheet [3]. The proportion who flow from each pool to the next is otherwise represented by a question mark, as these numbers are not represented by robust data.

We here present HBV within the framework for neglected tropical diseases (NTDs) [4], in order to highlight the ways in which HBV meets NTD criteria and to discuss the ways in which the NTD management paradigm could be used to strengthen a unified global approach to HBV elimination [5]. The major burden of morbidity and mortality from HBV is now borne by tropical and subtropical countries [6]. We here focus particular attention on Africa, as many African populations epitomize specific vulnerability to HBV [7]. However, the themes we represent are transferable to other low and middle-income settings, and are relevant on the global stage.

## CURRENT STRATEGIES FOR HBV CONTROL

Robust preventive vaccines have been rolled out in Africa since 1995 as a component of the Expanded Programme on Immunization (EPI). For adults with chronic infection and evidence of ongoing liver damage, a daily dose of suppressive antiviral therapy using nucleot(s)ide analogues (Table 1) successful at effecting viraemic suppression in the majority of cases, reducing complications and diminishing spread. Antiviral therapy does not commonly result in cure, due to the persistence of transcriptionally active DNA in the hepatocyte nucleus, but Interferon (IFN)-based therapy can increase rates of clearance.

**Table 1.**
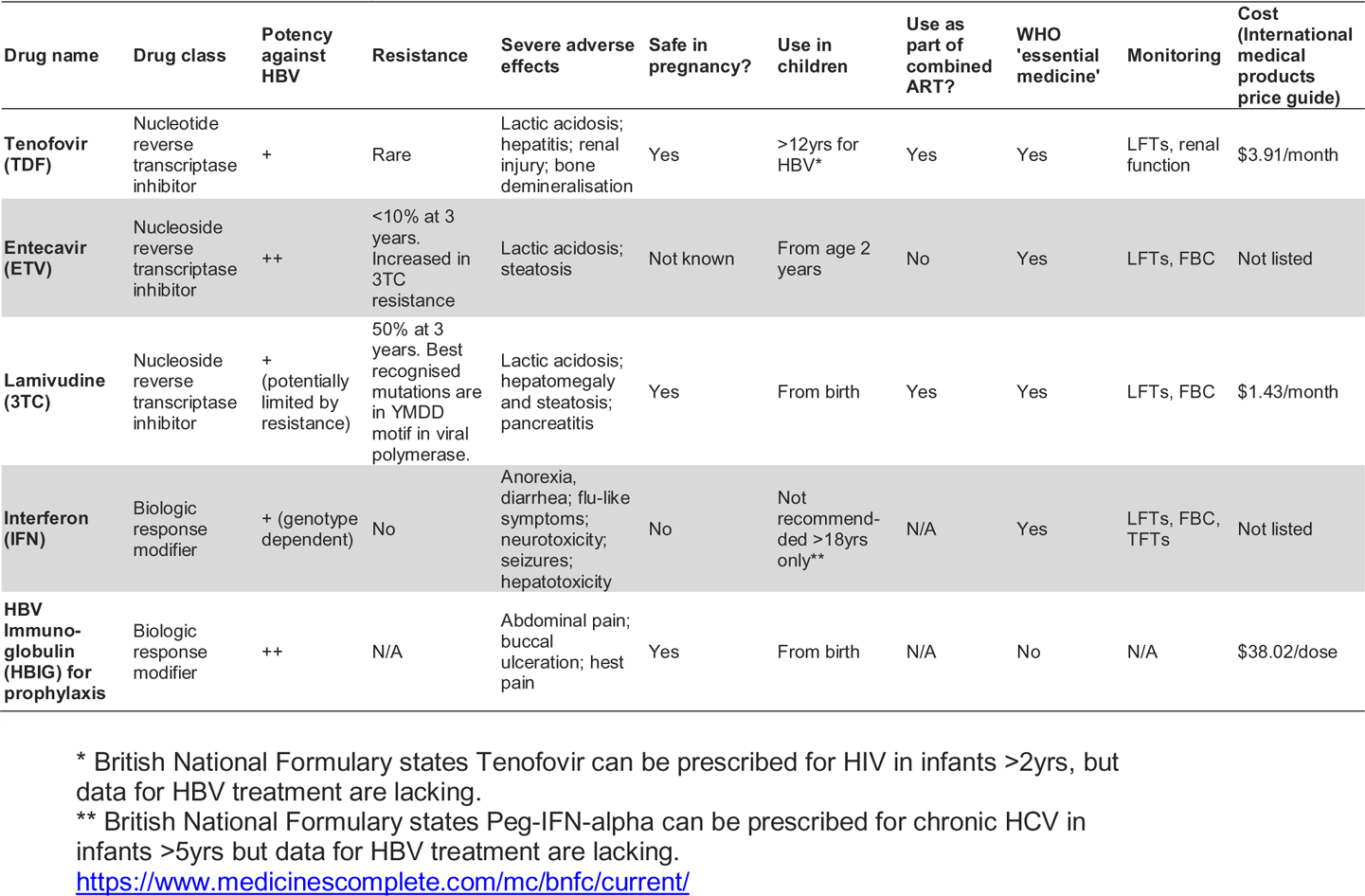
Drug therapy used to treat HBV. Costing is based on International medical products price guide: http://mshpriceguide.org/en (price for 3TC - South Africa Department of Health; price for TDF - Supply Chain Management Project; price for HBIG – Sudan MSF). WHO essential medicines: http://who.int/medicines/publications/essentialmedicines/EML_2015_FINAL_97amended_NOV2015.pdf?ua=1.

| Drug name | Drug class | Potency against HBV | Resistance | Severe adverse effects | Safe in pregnancy? | Use in children | Use as part of combined ART? | WHO 'essential medicine' | Monitoring | Cost (International medical products price guide) |
| --- | --- | --- | --- | --- | --- | --- | --- | --- | --- | --- |
| **Tenofovir (TDF)** | Nucleotide reverse transcriptase inhibitor | + | Rare | Lactic acidosis; hepatitis; renal injury; bone demineralisation | Yes | >12yrs for HBV* | Yes | Yes | LFTs, renal function | \$3.91/month |
| **Entecavir (ETV)** | Nucleoside reverse transcriptase inhibitor | ++ | <10% at 3 years. Increased in 3TC resistance | Lactic acidosis; steatosis | Not known | From age 2 years | No | Yes | LFTs, FBC | Not listed |
| **Lamivudine (3TC)** | Nucleoside reverse transcriptase inhibitor | + (potentially limited by resistance) | 50% at 3 years. Best recognised mutations are in YMDD motif in viral polymerase. | Lactic acidosis; hepatomegaly and steatosis; pancreatitis | Yes | From birth | Yes | Yes | LFTs, FBC | \$1.43/month |
| **Interferon (IFN)** | Biologic response modifier | + (genotype dependent) | No | Anorexia, diarrhea; flu-like symptoms; neurotoxicity; seizures; hepatotoxicity | No | Not recommended >18yrs only** | N/A | Yes | LFTs, FBC, TFTs | Not listed |
| **HBV Immuno-globulin (HBIG) for prophylaxis** | Biologic response modifier | ++ | N/A | Abdominal pain; buccal ulceration; hest pain | Yes | From birth | N/A | No | N/A | \$38.02/dose |

\* British National Formulary states Tenofovir can be prescribed for HIV in infants >2yrs, but data for HBV treatment are lacking.
\*\* British National Formulary states Peg-IFN-alpha can be prescribed for chronic HCV in infants >5yrs but data for HBV treatment are lacking. https://www.medicinescomplete.com/mc/bnfc/current/

Prevention of mother to child transmission (PMTCT) can be improved through a combination of routine antenatal screening, antiviral drugs during the latter stages of pregnancy, and HBV vaccination to the baby starting at birth. Where resources permit, HBV immunoglobulin (HBIG) can further reduce the risk of vertical transmission.

Despite the efficacy of these strategies in managing or preventing individual cases, these interventions do not currently offer a route to global HBV eradication, due to a shortage of investment and resources, the large pool of undiagnosed cases, lack of routine diagnostic screening, the high cost of IFN and HBIG, the lack of a curative therapy, substantial gaps in drug and vaccine coverage, and the potential for increasing drug resistance [8].

## APPLICATION OF NTD CRITERIA TO HBV

We have applied the WHO criteria for NTDs to HBV [4], and refer to case studies and experience from our own clinical practice (Suppl. data file) to illustrate how HBV in Africa fulfills NTD criteria.

### (i) NTDs ‘primarily affect populations living in tropical and sub-tropical areas’

Although HBV is endemic globally, the bulk of morbidity and mortality is now borne by low/middle income countries in tropical and sub-tropical regions [6, 9]. In Africa, many populations are particularly vulnerable due to co-endemic HIV infection and other co-infections, host and viral genetic factors, poverty, and lack of education and infrastructure [7]. In this setting, HBV has been eclipsed by the more acute and tangible health crisis of human immunodeficiency virus (HIV); only now in the ART era is it re-emerging as a visible threat [S2]. One illustration of this shift is the increase in deaths from HBV-related liver cancer over time that contrasts a reduction in AIDS deaths [10].

### (ii) NTDs ‘disproportionately affect populations living in poverty; and cause…. morbidity and mortality, including stigma and discrimination’

HBV is part of a cycle of poverty, with a high burden of morbidity and mortality in young adults [S1, S4, S9]. The economic burden on individual families can be particularly catastrophic in low and middle income settings [11] [S4, S5, S7], although robust data are lacking for Africa. In resource-poor settings, lack of education and scarce healthcare resources impinge on successful diagnosis and monitoring [S4, S7], as well as failure to control symptoms where relevant [S9]. Stigma and discrimination are often invisible, but can be potent and highly relevant challenges to the success of scaling up interventions for prevention, diagnosis, and treatment [12] [S5, S6, S7].

### (iii) NTDs are ‘immediately amenable to broad control, elimination or eradication by applying… public health strategies’

We already have an armamentarium of strategies with which to tackle HBV prevention and treatment (Figure 2). In order to be widely and robustly deployed, these approaches should interlink with existing resources and infrastructure wherever possible [S2].

**Figure 2.**
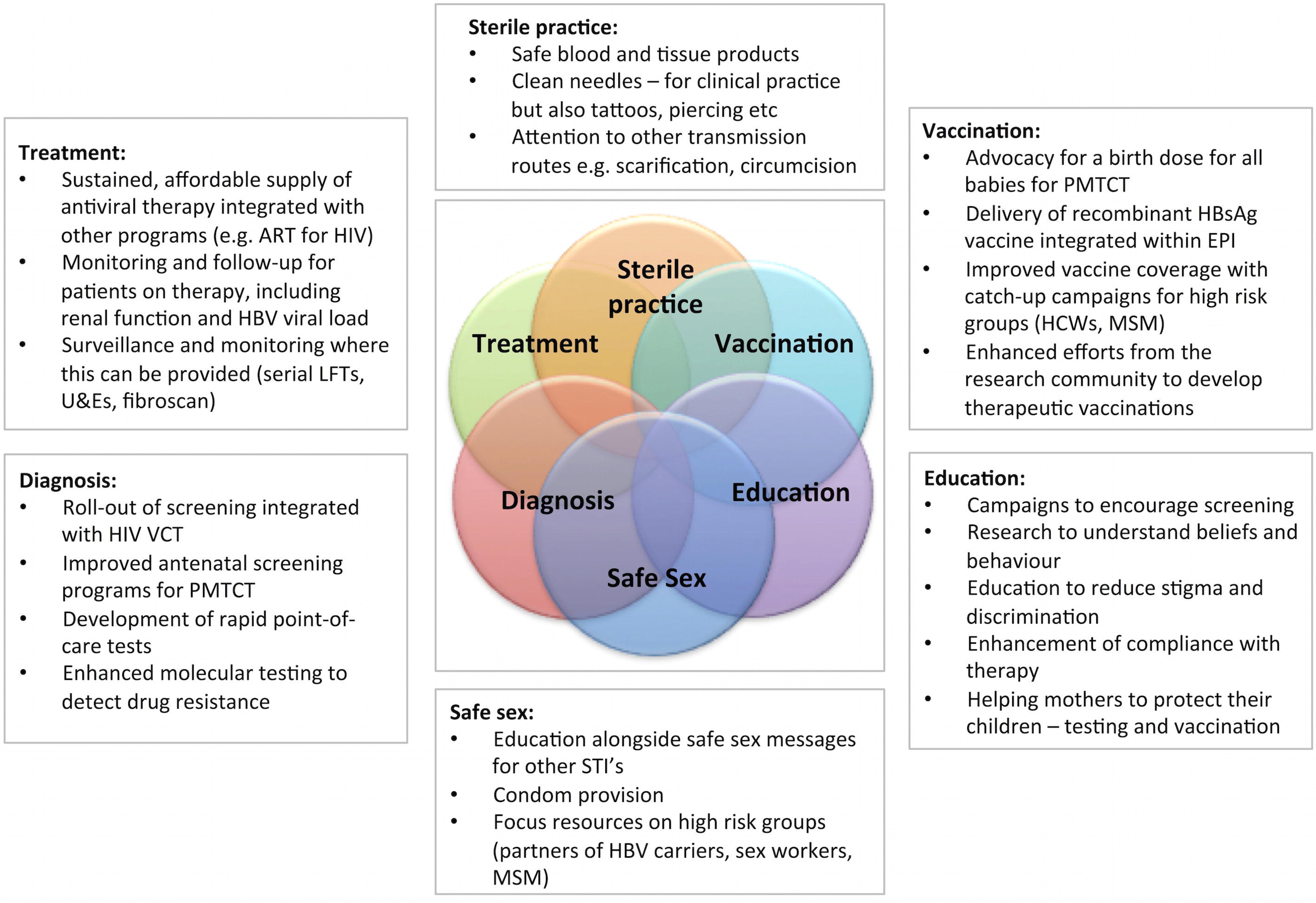
A package of interventions to move towards elimination of HBV infection as a public health threat. Suggested measures are aligned with WHO interventions for NTDs.

### (iv) NTDs are ‘relatively neglected by research – i.e., resource allocation is not commensurate with the magnitude of the problem’

Compared with other blood-borne viruses, namely HIV and hepatitis C virus, which infect substantially lower numbers [7], HBV has attracted far fewer research resources, and this gap may actually be widening over time [13]. HBV mortality (887,000 deaths / year [3]) is now twice that of malaria (429,000 deaths / year [14]) but, malaria receives nearly five-fold more funding (Figure 3). Moreover, development of clinical programs for hepatitis testing and treatment are fragmented in comparison to the progressive infrastructure that has emerged to tackle HIV [S7].

**Figure 3.**
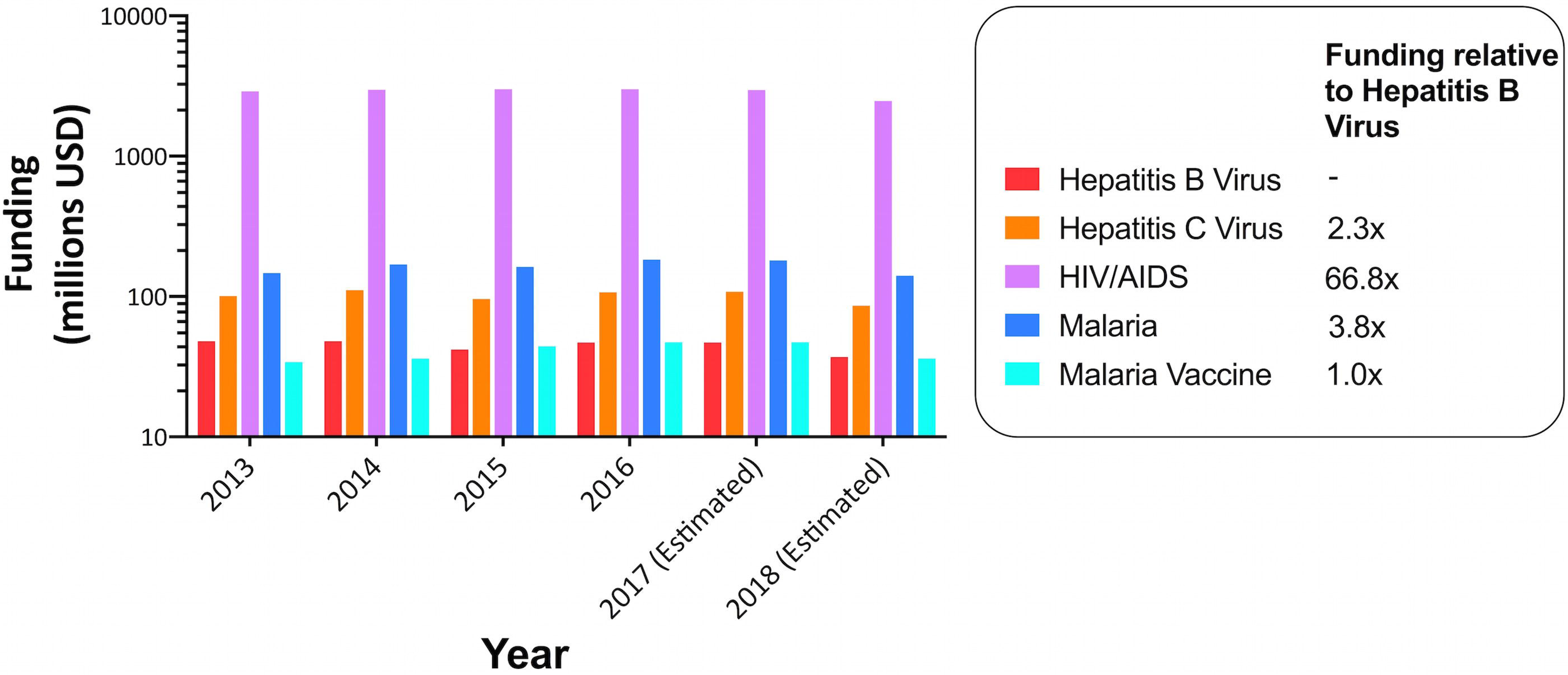
Funding allocations for HBV, HCV, HIV and malaria, 2013-2018. Data from the US National Institutes for Health (NIH) estimated funding for research, condition and disease categories 2013-2018 (projected), available at https://report.nih.gov/categorical_spending.aspx. Figures for the projected funding allocation (for 2018) relative to HBV are given. *Research into ‘malaria’ and ‘malaria vaccine’ have been subdivided in the dataset.

## RECOMMENDATIONS BASED ON NTD FRAMEWORK

Even for an organism that is not officially recognized as an NTD, there is much to be learnt from the NTD paradigm that could accelerate progress in tackling HBV. The ethos of combining several public health strategies, and integrating care for different diseases, is captured by the approach advocated for NTDs [4], and is also a helpful model for HBV. Particularly in the African subcontinent, where other NTD models have had significant impact [15], using this framework for HBV could promote awareness, leverage advocacy and resources, and promote integration of HBV prevention and treatment into existing HIV infrastructure [5].

In the following section, we use suggested interventions for NTDs to discuss briefly how these are pertinent to reducing – and ultimately eliminating – HBV infection as a public health threat.

### (i) ‘Intensified case management’

Based on the significant numbers of individuals lost at every step of the ‘cascade’ from diagnosis through to successful treatment and prevention (Figure 1), enhanced efforts are needed to promote linkage through care pathways. Enhanced HBV testing is crucial to facilitate entry into clinical care, allowing treatment to reduce the risk of onward spread, including underpinning PMTCT [S8]. Initially, this may rely on using existing diagnostic platforms (based on serology), but investment is required in developing and rolling out new approaches, including molecular testing strategies that are more sensitive, provide enhanced data (e.g. detection of drug resistance), and are fast enough to enable point-of-care testing. This can often be transferred from technology that has been initially developed for the diagnosis of other diseases.

The role and significance of stigma associated with HBV infection in Africa is largely unreported in the literature. However, individual testimony leaves no doubt that this is a significant barrier to diagnosis and clinical care [S5, S6]. Gaining a better understanding of the extent and nature of stigma and discrimination in different populations is a crucial first step, in parallel with enhanced efforts to educate patients, health care workers and the public.

### (ii) ‘Preventive chemotherapy’

Although antiviral therapy for HBV is generally regarded as treatment rather than prevention, in the majority of cases it renders individuals aviraemic, preventing onward transmission. Antiviral therapy for HBV (Table 1) should be made accessible, ideally capitalizing on the supply chains and distribution infrastructure that have been developed for HIV (and/or other prevalent infections, such as tuberculosis and malaria) [5]. Research efforts are still required to identify prognostic factors that predict differential response to therapy and allow tailoring of care.

PMTCT can progressively become a realistic goal by expanding access to antenatal diagnostics, simple treatment interventions such as maternal tenofovir during trimester three, and HBV vaccination for all babies, with the first dose delivered at birth [8] [S8]. Vaccination remains a cornerstone of prevention, but more work is needed to investigate the most effective catch-up immunization strategies to reduce the burden of HBV infection at a population level [S3, S4].

### (iii) ‘Sanitation and hygiene’

Although this category of interventions is conventionally applied to reducing food and water-borne infections, we here broaden our interpretation to include other aspects of prophylaxis. Safety and security of medical supplies has increasingly improved to reduce nosocomial transmission of blood borne viruses over recent decades [S3]. However, sterile practices need to be more widely promoted and guaranteed, to assure the safety of other procedures such as scarification, tattoos, piercings and circumcision that may occur in community settings. Provision of condoms alongside education regarding safe sex, particularly for high risk groups such as sex workers and men who have sex with men, is another important strategy for prevention.

## CONCLUSIONS

Elimination of HBV infection has gained status within international health and development agendas, but is a complex clinical and public health challenge that currently lacks proportionate multi-lateral commitment from pharma, government, commissioners, funders and the research community. The many parallels with other NTDs are clearly exemplified by vulnerable populations of the African subcontinent. By viewing HBV within the NTD framework, we can improve approaches to reducing the burden of disease and move towards eventual elimination.

## SUPPORTING INFORMATION LEGEND

This document contains supplementary data to support our view that Hepatitis B Virus (HBV) can helpfully be represented within the framework set out for Neglected Tropical Diseases by the World Health Organization (WHO) [1]. This is in line with aims stated within Sustainable Development Goals [2]. Complementary evidence gathered from different locations in Africa illustrates the ways in which HBV infection meets the criteria for NTDs. These scenarios (labelled S1 to S9, and presented by geography from South to North) contribute important insights into how the NTD paradigm can be helpful in informing strategies to improve diagnosis, treatment and prevention of HBV infection, with the ultimate goal of eliminating infection as a public health threat.

